# EcoEvoApps: Interactive Apps for Theoretical Models in Ecology and Evolutionary Biology

**DOI:** 10.1101/2021.06.18.449026

**Authors:** Rosa M. McGuire, Kenji T. Hayashi, Xinyi Yan, Marcel Caritá Vaz, Damla Cinoğlu, Madeline C. Cowen, Alejandra Martínez-Blancas, Lauren L. Sullivan, Sheila Vazquez-Morales, Gaurav S. Kandlikar

**Affiliations:** Department of Ecology and Evolutionary Biology, University of California, Los Angeles; Department of Integrative Biology, University of Texas at Austin; Institute for Environmental Science and Sustainability, Wilkes University; Departamento de Ecología y Recursos Naturales, Facultad de Ciencias, Universidad Nacional Autónoma de México; Division of Biological Sciences, University of Missouri, Columbia; Division of Plant Sciences & Technology, University of Missouri, Columbia

**Keywords:** mathematical modeling, R package, shiny apps, ecological theory, teaching

## Abstract

1. The integration of theory and data drives progress in science, but a persistent barrier to such integration in ecology and evolutionary biology (EEB) is that theory is often developed and expressed in the form of mathematical models that can feel daunting and inaccessible for students and empiricists with variable quantitative training and attitudes towards math.
2. A promising way to make mathematical models more approachable is to embed them into interactive tools with which one can visually evaluate model structures and directly explore model outcomes through simulation.
3. To promote such interactive learning of quantitative models, we developed EcoEvoApps, a collection of free, open-source (R/Shiny) apps that include model overviews, interactive model simulations, and code to implement these models directly in R. The package currently focuses on canonical models of population dynamics, species interaction, and landscape ecology. We also outline a vision and approach for growing the collection to include more models from across EEB.
4. These apps help illustrate fundamental results from theoretical ecology and can serve as valuable teaching tools in classroom settings. We present data from student surveys which show that students rate these apps as useful learning tools, and that using interactive apps leads to substantial gains in students’ interest and confidence in mathematical models. This points to the potential for interactive activities to make theoretical models more accessible to a wider audience, and thus facilitate the feedback between theory and data across ecology and evolutionary biology.

## Introduction

Integrating theory with insights from observations and experiments is a fundamental driver of progress across the life sciences (Jungck 1997, Shou et al. 2015), including in ecology and evolutionary biology (EEB) (Marquet et al. 2014, Servedio et al. 2014). While not all theory is mathematical, research that synthesizes data with mathematical models can enable generalization across systems, promote a deeper conceptual understanding of biological systems by clarifying the role and consequences of different biological factors, help disentangle complex interactions and feedbacks, and highlight important areas for further study (Haldane 1964, Caswell 1988). Such integration can also have important applications in biological forecasts and in informing actions and policies at the interface of science and society (Conway 1977, Wainwright et al. 2018). Despite widespread agreement between empiricists and theoreticians that more synergism between these two approaches towards EEB research can yield fruitful insights (Jeltsch et al. 2013, Scheiner 2013, Haller 2014, Shou et al. 2015), there are numerous barriers that limit such integration.

One such barrier towards more integration is that the language of mathematical models and their analytical solutions may seem foreign to those who come to EEB from a more empirical background. As a result, equation-heavy papers tend to be cited less often (Fawcett and Higginson 2012), and instructors of quantitative courses tend to receive worse student evaluations than those who de-emphasize quantitative topics (Uttl et al. 2013, Kreitzer and Sweet-Cushman 2021). However, while many authors have called for an increased emphasis on quantitative training at all stages in EEB education, these calls focus primarily on an increased emphasis on statistical models (e.g. Ellison and Dennis 2010) or on programming/computational skills (e.g. Losos et al. 2013, Feng et al. 2020), with relatively few advances in the pedagogy of theoretical models (but see Lehman et al. 2020, Grainger et al. 2022). Across quantitative biology more broadly, a growing body of research suggests that interactive tools that allow users to independently explore model structure and outcomes help increase student interest and understanding of quantitative concepts (e.g. Thompson et al. 2010, Feser et al. 2013, Ou et al. 2022). Establishing a platform for interactive simulations of EEB models thus has the potential to facilitate communication and collaboration between theoretical and empirical researchers.

Here we describe EcoEvoApps, an open-source R package (ecoevoapps) and website (https://ecoevoapps.gitlab.io) that provides a collection of freely available interactive apps that simulate fundamental EEB models. The package also includes functions to directly run models through the R console, and can thus serve as a bridge to help users become familiar with coding and implementing theoretical models. We illustrate how these apps can be used to help communicate and learn insights from theoretical models, both at the level of an individual seeking to gain more familiarity with a model, and in large undergraduate classroom settings. We actively invite anyone who wishes to contribute to the project by writing new apps, reviewing and/or adding new features to existing apps, translating apps into other languages, or contributing teaching plans, to join our community.

## Package overview

### Interactive (Shiny) apps

At the heart of ecoevoapps are 11 interactive apps (Table 1), which we expect to be the primary avenue through which most users interact with the package. We chose the models to include in this first release of ecoevoapps by surveying syllabi for undergraduate ecology courses and commonly-used textbooks (Gotelli 2008, Begon and Townsend 2020). We expect to build on this collection with future releases of the package. Some apps implement the dynamics of one specific model (e.g. the abiotic resource competition app, which models two species competing for two essential resources (Tilman 1980)), while other apps present several closely related models. For example, the predator-prey dynamics app includes a tab that presents the classic Lotka-Volterra model, and other tabs with model extensions that integrate logistic growth in the prey and/or a type II functional response for the predator (Fig. 1). Each app includes a brief description of the model structure and history, a table with parameter definitions, and references to relevant literature. A core set of nine apps are available in English, Spanish, and Chinese, Turkish, and Portuguese (Table 1). We plan to continue adding new apps and translating existing apps both internally and by soliciting contributions from community members (see “Contributing to EcoEvoApps” below).

**Table 1:** Models and functions included in the `ecoevoapps` package. In addition to the functions listed in the table, the package also includes 11 functions with the prefix shiny_ that can be used to deploy shiny apps directly from the command line (https://ecoevoapps.gitlab.io/docs/reference/index.html#run-shiny-apps).

| Model | Link to shiny app | Functions for running the models/shiny apps | Functions for plotting model outputs |
| --- | --- | --- | --- |
| Population dynamics in continuous time | 中文;<br>Español;<br>English;<br>português;<br>Turkish | run_exponential_model()<br>run_logistic_model()<br>shiny_singlepop_continuous() | plot_continuous_population_growth() |
| Population dynamics in discrete time | 中文;<br>Español;<br>English;<br>português;<br>Turkish | run_discrete_exponential_model()<br>run_discrete_logistic_model()<br>run_beverton_holt_model()<br>run_ricker_model()<br>shiny_population_growth_discrete() | plot_discrete_population_growth()<br>plot_discrete_population_cobweb() |
| Structured population growth | 中文;<br>Español;<br>English;<br>português;<br>Turkish | run_structured_population_simulation()<br>shiny_structured_population() | plot_leslie_diagram()<br>plot_structured_population_size()<br>plot_structured_population_lambda()<br>plot_structured_population_agedist() |
| Lotka-Volterra competition | 中文;<br>Español;<br>English;<br>português;<br>Turkish | run_lvcomp_model()<br>shiny_lvcomp_model() | plot_lvcomp_time()<br>plot_lvcomp_portrait() |
| Predator-prey dynamics | 中文;<br>Español;<br>English;<br>português;<br>Turkish | run_predprey_model()<br>shiny_predprey() | plot_predprey_time()<br>plot_predprey_portrait() |
| Competition for abiotic resources | 中文;<br>Español;<br>English;<br>português;<br>Turkish | run_abiotic_comp_model()<br>run_abiotic_comp_rstar()<br>shiny_abiotic_comp() | plot_abiotic_comp_time()<br>plot_abiotic_comp_portrait() |
| Competition for a biotic resource | 中文;<br>Español;<br>English;<br>português;<br>Turkish | run_biotic_comp_model()<br>shiny_biotic_comp() | plot_biotic_comp_time()<br>plot_functional_responses() |
| Compartment models of infectious disease dynamics | 中文;<br>Español;<br>English;<br>português;<br>Turkish | run_infectiousdisease_model()<br>shiny_infectious_disease() | plot_infectiousdisease_time()<br>plot_infectiousdisease_portrait() |
| Island biogeography | 中文;<br>Español;<br>English;<br>português;<br>Turkish | run_ibiogeo_model()<br>shiny_ibiogeo_model() | none (run_ibiogeo_model() itself returns plots) |
| Smith-Fretwell model | English | run_smithfretwell_model()<br>shiny_smith_fretwell() | plot_smithfretwell_model() |
| Metapopulation dynamics | English | run_source_sink()<br>shiny_source_sink() | plot_source_sink() |

**Figure 1:**
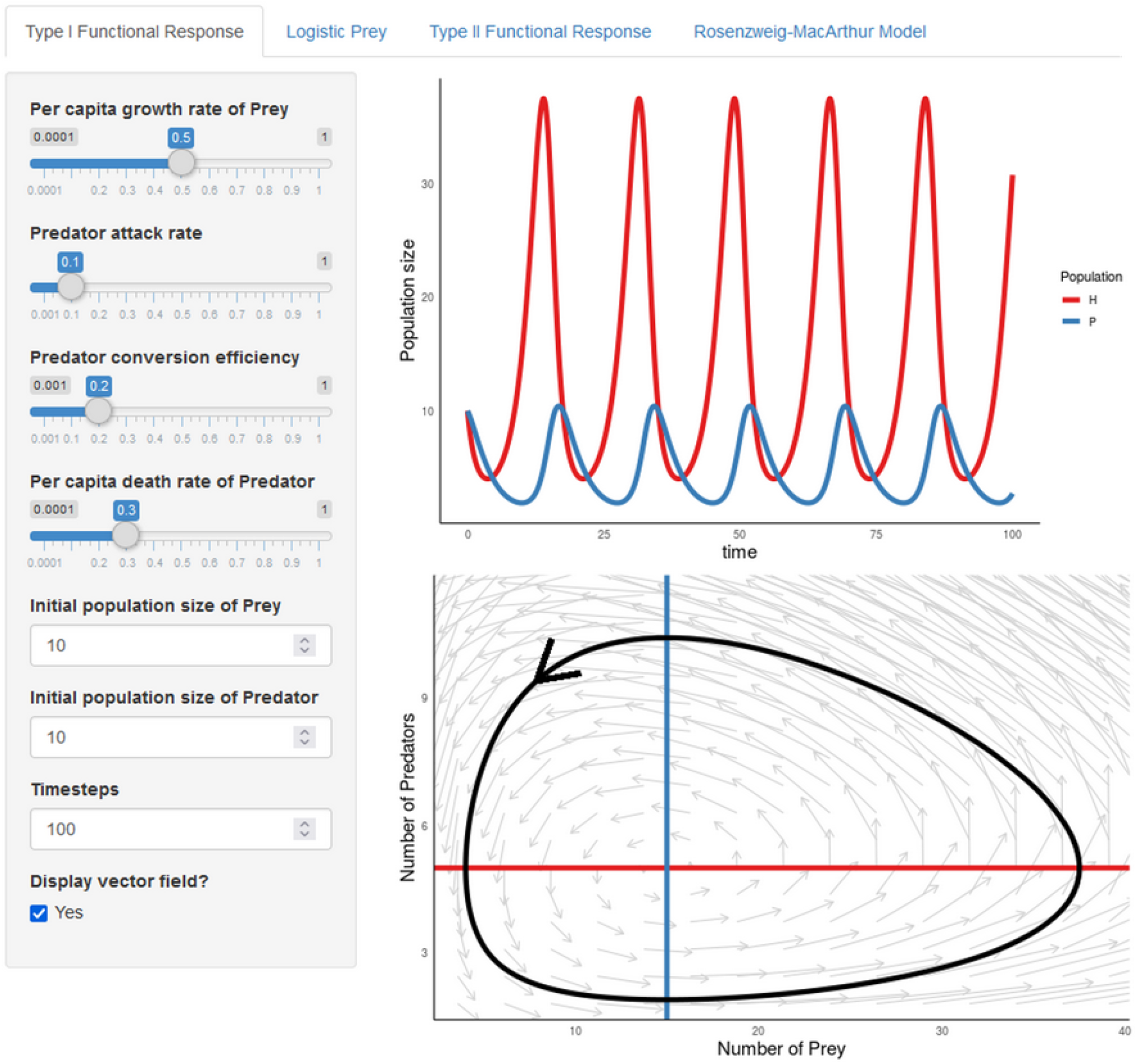
Screenshot of the predator-prey dynamics shiny app being used to simulate Lotka-Volterra dynamics. Users set parameter values on the left-hand panel, and these inputs are used to generate the population trajectory and phase portrait on the right. In addition to this interactive component, the shiny app also includes a verbal description of the model, the model equations, and a parameter table. The app also includes three other tabs that incorporate logistic growth in the prey, type II functional response for the predator, or both. The app is available online at https://ecoevoapps.shinyapps.io/predator_prey_dynamics/, or can be deployed from the R command line with `shiny_predprey()`.

The shiny apps are freely available online on RStudio’s shinyapps.io servers (links available in Table 1), or can be launched locally from users’ personal computers from the R console. For such deployment, the package provides a series of functions with the prefix shiny_ that launch the apps. The package also includes a vignette with instructions for users who wish to customize and deploy their own instance of an app, e.g. for hosting on institutional servers or to modify an app’s content for a specific classroom lesson. Finally, the package also includes model-specific vignettes with instructions for simulating the model dynamics directly through `R (e.g. vignette(“predator-prey-interactions”))`.

### Functions for simulating and visualizing model dynamics

Under the hood, the shiny apps use functions in the ecoevoapps package to simulate and visualize model dynamics (Table 1). Simulations are conducted by functions with the prefix `run`_, which take as their input the parameter values and other relevant information for the particular model. For example, `run_predprey_model()` requires as inputs a vector defining the parameter values (`params`), a vector of the initial population sizes for the predator and prey species (`init`), and the time steps over which to run the model (`time`). The function returns a dataframe of the population sizes for each species over the specified time series. The package also includes a series of plotting functions prefixed `plot_`, which take as their input the object returned by the corresponding run_ function, and in turn return a `ggplot2` object. Using `ecoevoapps` functions, the outputs in Fig. 1 can be generated with the following code:

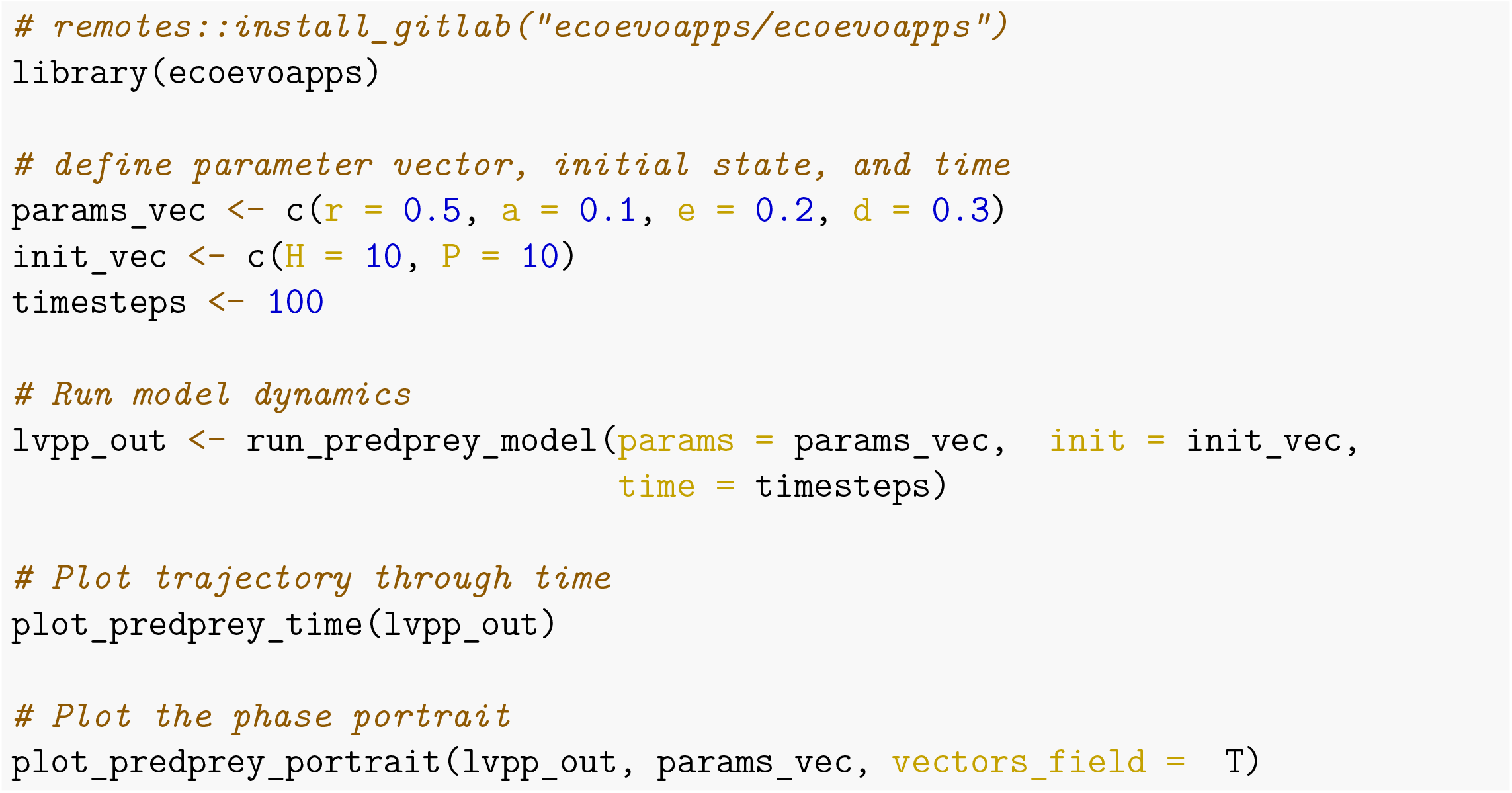

The complete list of functions for simulating and plotting model dynamics is provided in Table 1. Each function’s usage is documented in the package, and suites of functions relevant to different models are described in the corresponding vignettes. While we expect the interactive apps to be the primary mode for most user’s engagement with the package, users familiar with R — or those who wish to build this familiarity — can use these functions to conduct visualizations or analyses beyond those presented in the apps. Thus, the package can also serve as a gateway for users to implement and manipulate mathematical models at the command line.

### Installation and dependencies

The `ecoevoapps` package can be installed from GitLab: `remotes::install_gitlab(“ecoevoapps/ecoevoapps”)`. The package depends on functions from `deSolve` (Soetaert et al. 2010), diagram (Soetaert 2020), patchwork (Pedersen 2020), and various packages within the tidyverse (Wickham et al. 2019). We have tested the `ecoevoapps` package on `R` versions >4, and have tested the shiny apps on Firefox, Chrome, and Safari.

### Contributing to EcoEvoApps

This manuscript describes the first release of EcoEvoApps, and we envision this package to grow as a collaborative and inclusive effort. In particular, our overarching goal is to leverage the diverse expertise of the EEB community to build an open educational resource that facilitates dialogue between theoretical and empirical research. As such, EcoEvoApps offers several mechanisms by which educators, researchers, and students can contribute to the project. These mechanisms include (1) writing and contributing new apps, (2) revising existing apps, (3) providing feedback, translating apps, or requesting new apps or features, and (4) contributing classroom activities or other use-cases involving the use of one or more of the apps. Detailed contribution guidelines are provided as a vignette (`vignette(“contributing”`)). Contributors are acknowledged in the package source code, as well as on the project homepage (https://ecoevoapps.gitlab.io/people/).

## Use cases

### Communicating and learning insights from classic models

Theoreticians and empiricists alike can use shiny apps to help communicate and learn insights from mathematical models. For example, the paradox of enrichment (Rosenzweig 1971) can be visualized with the predator-prey model app by altering the value of the prey carrying capacity (*K*) in the *MacArthur-Rosenzweig model* tab. Low values of prey carrying capacity result in a stationary equilibrium or one with stable oscillations, while high values of prey carrying capacity — as might occur when a system is “enriched” — result in unstable oscillations that ultimately limit the system’s persistence. In particular, careful exploration of the parameter *K* can reveal the logic behind Rosenzweig (1971)’s conclusion that the system can persist with stable oscillations or a stationary equilibrium only when the equilibrium point (intersection of the two isoclines) occurs to the right of the hump in the prey isocline (Fig. 2).

**Figure 2:**
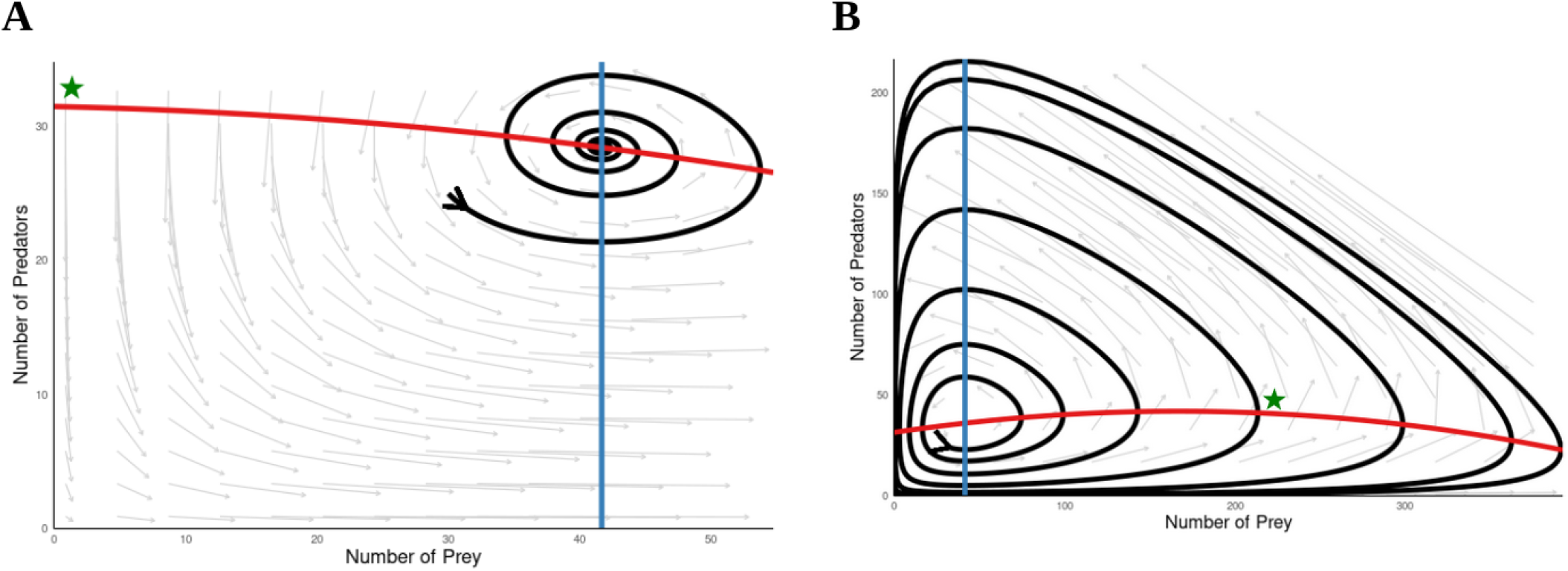
Screenshots of the predator-prey dynamics shiny app being used to simulate the Macarthur-Rosenzweig model. Panel A shows dampened oscillations arising at low prey carrying capacities (K = 200) when the predator (blue) isocline intersects the prey (red) isocline to the right of the “hump” (indicated by the green star, which was added onto the screenshot). In contrast, panel B shows the unstable oscillations that arise under high prey carrying capacity (K = 500) when predator isocline intersects the prey isocline to the left of the hump (green star).

### Classroom teaching

ecoevoapps can also be used as a formal instruction tool for teaching mathematical models. To evaluate the value of these apps in classroom settings, we surveyed 51 students who used the shiny apps for Island Biogeography and Lotka-Volterra competition to learn these topics in an upper-division Ecology course at the University of California, Los Angeles (UCLA, see supplements S1-S5 for details). The learning activity included short (∼15 minutes) video lectures that presented an overview of the model and the shiny app, followed by a worksheet that navigated students through a guided exploration of the model (worksheet available in Supplement S3). After completing the activity, students rated on a scale of 1-7 the degree to which the apps helped them understand the model as a whole, as well as specific topics associated with the model. An overwhelming majority of students (40/51) reported that the apps were moderately to very helpful for learning the models as whole (response of 6 or 7, Fig. 3A). The apps also appear to help students better understand specific ideas related to the models (e.g. students report that they better understand the concepts of “carrying capacity” or “coexistence” after using the Lotka-Volterra competition app, Fig. 3B). We also conducted similar surveys of students in a General Ecology course at the University of Missouri (MU), with similar results (Supplement S2). In particular, by tracking individual students’ interest and confidence in models before and after the activity, we found that using interactive apps led to substantial gains in student confidence, especially among students who express higher interest in related topics (Fig. S2.1). Classroom surveys were reviewed by the UCLA Institutional Review Board and MU Institutional Review Board were determined to constitute “exempt” studies (UCLA IRB #20-002179; MU IRB Project #2031063, Review #276104).

**Figure 3:**
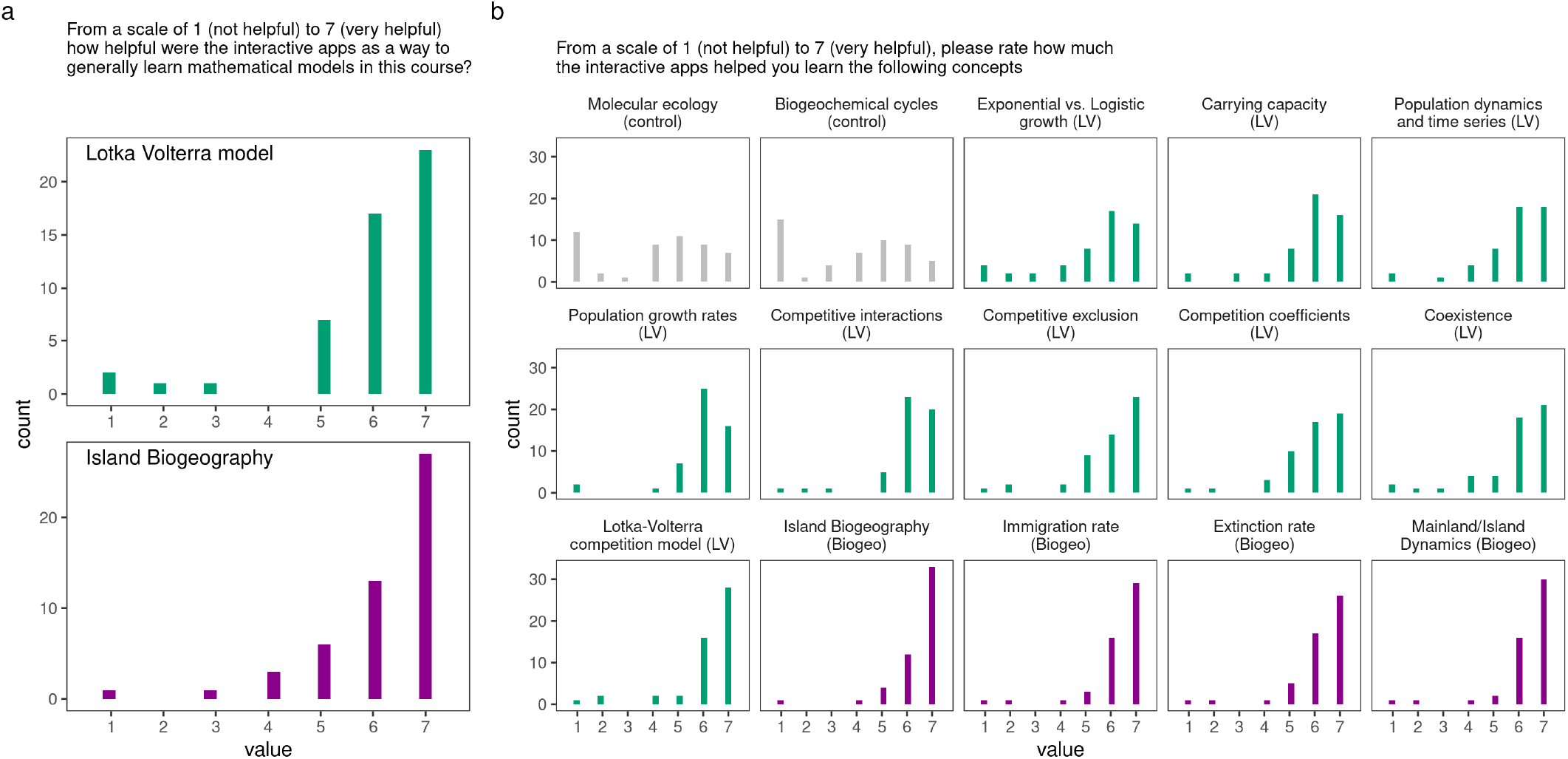
Students at UCLA (n = 51) generally rated the Lotka-Volterra competition and Island Biogeography apps to be valuable tools to help learn the models overall (a), as well as for specific topics within each model (b). Green histograms indicate topics related to the Lotka-Volterra competition model, purple histograms indicate topics related to Island Biogeography, and grey histograms indicate topics unrelated to either activity, which served as a control.

## Conclusions and outlook

Integrating theoretical and empirical approaches is often heralded as an ideal path for progress in ecology and evolutionary biology (Jeltsch et al. 2013, Shou et al. 2015, Laubmeier et al. 2020, Servedio 2020), but such integration remains relatively limited (Scheiner 2013). One likely barrier is that students are often not exposed to extensive quantitative training in traditional biology curricula (Chiel et al. 2010), and as a result, theoretical models remain intimidating for many empirical researchers (Haller 2014, Grainger et al. 2022). While simulation-based learning may not provide all the same insights as analytical solutions, platforms like `R` and shiny allow us to build tools that give everyone easier access to theoretical insights can otherwise take years of quantitative training to grasp. We leveraged these advances to build EcoEvoApps, a collection of web apps that allow users to interactively explore theoretical models, adding to a variety of existing interactive EEB education web resources (e.g. Evo-Ed (http://www.evo-ed.org), HHMI BioInteractive (https://www.biointeractive.org), Populus (Alstad 2001)). A key distinguishing feature is that unlike these other resources, ecoevoapps is entirely open-source and written in R. As such, it is easily accessible and customizable by others in the EEB community, where R is among the most commonly used programming languages (Gentleman et al. 2004, Lai et al. 2019). Moving forward, we will prioritize incorporating mathematical models from evolutionary biology and population genetics into the package to complement the current ecological focus. Building on our preliminary evidence that shiny apps are useful tools for teaching quantitative models in classroom settings, we also plan to develop and evaluate new lesson plans for EEB educators teaching mathematical models.

## Acknowledgments

We acknowledge the Gabrielino/Tongva peoples as the traditional land caretakers of Tovaangar (the Los Angeles Basin and Southern Channel Islands), where the University of California, Los Angeles (UCLA) is located. We also respectfully acknowledge that the University of Missouri is located on the traditional, ancestral lands of the Osage, Omaha, and Kaw peoples, among others. We thank Gary Bucciarelli for helping conduct surveys at UCLA, and Mayda Nathan, Chris Muir, and Daniel Gruner for contributing apps to the EcoEvoApps website. We thank Caroline Farrior for comments on the manuscript. This material is based upon work supported by the National Science Foundation Graduate Research Fellowship Program to RMM under Grant No. (DGE-2034835). RMM was also supported by the Eugene V. Cota-Robles Fellowship. MCV was supported by a CAPES doctoral fellowship (2014, BEX:10079-13-0). MCC was supported by an NIH Systems in Integrative Biology Training Grant. Any opinions, findings, and conclusions or recommendations expressed in this material are those of the author(s) and do not necessarily reflect the views of the National Science Foundation or the NIH.

## Notes

https://ecoevoapps.gitlab.io/

### Competing Interest Statement

The authors have declared no competing interest.

### Summary of Updates

Restructured the manuscript -- substantially simplifies the classroom data component, and now focus more on the actual implementation of the package. The package now also includes all focal apps in 5 languages, with many translations done by newly added authors.

