## Supplements 1-5 for "EcoEvoApps: Interactive Apps for Theoretical Models in Ecology and Evolutionary Biology"

**Last rendered**: 25 May 2022

#### Supplement S1: Classroom survey methods

To determine the value of apps in `ecoevoapps` for teaching theoretical ecology concepts, we surveyed students who used EcoEvoApps in two upper-division undergraduate ecology courses. Students in EE BIOL 122 (Ecology) at the University of California, Los Angeles ("UCLA") completed activities using the Lotka-Volterra Competition and Island Biogeography apps, while students in BIOL 3650 (General Ecology) at the University of Missouri, Columbia ("MU") completed activities using the Lotka-Volterra Competition and Infectious Disease apps, and students. Students were offered extra-credit points for completing a learning activity (Supplement S4) using the interactive apps and were encouraged to complete a survey (Supplement S5) before and after completing the activity. Classroom research was reviewed by the MU Institutional Review Board (Project #2031063; Review #276104) and the UCLA Institutional Review Board (IRB#20-002179), and was determined to constitute "exempt" studies.

The pre-activity survey at both universities asked students to rate their interest in six ecological sub-disciplines (population ecology, community ecology, conservation ecology, ecosystem ecology, global change ecology, and disease ecology). Students were also asked to report their confidence on a scale of 1-7 in specific concepts that were either relevant to the focal learning activity, or control topics that were related to a different topic covered in the course (Biogeochemistry/Molecular ecology in the UCLA class, Structured population growth in the MU class). The learning activities at both universities were worksheets that guided students through manipulating the apps with various parameter combinations, which were chosen to illustrate specific biological scenarios. The UCLA learning activity also included a short instructional video explaining the conceptual basis of the focal model, as well as a demonstration of the interactive app. Worksheets from both classes are available in Supplement S2 below.

At UCLA, the post-activity survey asked students to rate on a scale of 1-7 how helpful they found the interactive apps as a way to learn a series of relevant concepts, and also requested general feedback to help improve the apps through a free-response question. Although this design did not allow us to track potential changes in individual students' confidence with the topics, we used these responses to evaluate whether students generally found the interactive apps to be a helpful and engaging way to learn ecological models. All survey questions are available in Supplement S3 below.

At MU, questions on the post-activity survey were the same as those on the pre-activity survey. This allowed us to calculate, for each student, the change in confidence in specific concepts related to the focal models before and after using an interactive app. We evaluated whether using the interactive apps changed students' self reported confidence between the pre- and post-activity surveys at MU by computing the normalized change metric (Marx and Cummings 2007), modified to reflect 7 as the maximum score for each question:

$$c = \begin{cases} \frac{\text{post-pre}}{7-\text{pre}} & \text{post} > \text{pre} \\ \text{drop} & \text{pre} = \text{post} = 7 \\ 0 & \text{post} = \text{pre} \\ \frac{\text{post-pre}}{\text{pre}} & \text{post} < \text{pre} \end{cases}$$

This metric scales each students' realized gains or losses relative to the maximum possible gain or loss according to the pre-activity survey, thus allowing for more fair comparisons among questions. Following Marx and Cummings (2007), we interpret the results in terms of the mean and standard error of the normalized change for each question. We used data from the UCLA surveys to evaluate whether students found the apps useful for learning various ecological concepts, and read through all feedback to identify the salient themes. All analyses were conducted in R version 4.0.5 (R Core Team 2021). R code to recreate analyses is provided in Supplement S6, and will be archived in an appropriate repository upon acceptance.

#### Supplement S2: Results from classroom surveys

Among students at UCLA, an overwhelming majority indicated that the interactive apps were a moderately helpful to very helpful way to learn the related models (See main text). The free response feedback about the apps from UCLA students largely fell into four themes: visualization, help understanding concepts, manipulating parameters/making connections, and applicability of the models (see Supplement S5 below). A vast majority of the students (88%) reported that the models helped them learn the overall models (Lotka-Volterra/Island Biogeography) or identified a specific concept that the apps helped them understand, such as coexistence and population dynamics. In particular, 24% of students highlighted that being able to manipulate individual parameters and observe model outcomes was helpful in understanding the mathematical basis of the models, which was difficult to grasp without directly interacting with the model. 22% of students mentioned the value of the visualizations generated by the apps (time series, isoclines) for understanding model outcomes. Finally, 6% of students reported that they better understood the models by working through the case studies presented in the worksheet. Most students left comments that integrated across each of these themes, e.g.:

> The Lotka-Volterra simulation helped me understand what the Lotka-Volterra [model] predicts because it was more hands on than listening to its explanation during lecture. The island [biogeography] simulation also made it easy to understand how different variables and values of size/distance affect island populations. Visualizing these concepts made the model very clear.

At MU, students reported substantial gains in their confidence in all concepts related to the Lotka-Volterra competition model after completing the related activity (mean $\pm$ SE of normalized change across all Lotka-Volterra-related concepts = 0.241 $\pm$ 0.0244; normalized change across all structured population growth concepts (control) = 0.152 $\pm$ 0.033, Fig. S2.1a). Students with higher levels of interest in community ecology generally reported higher gains in their confidence after using the app than students who were largely uninterested in community ecology (Fig. S2.1b). Within the Lotka-Volterra category, students reported highest gains in confidence for general concepts related to the model (e.g. positive or negative species interactions; two-species interactions) rather than for specific concepts or model parameters (e.g. competition coefficients; inter- vs.intra-specific interactions) (Fig. S2.1c).

Students also reported substantial gains in confidence in concepts related to the SIR disease dynamics model as a whole (mean SE of normalized change across all SIR-related concepts = 0.219 $\pm$ 0.0324; normalized change across all structured population growth concepts (control) = 0.164 $\pm$ 0.0329, Fig. S2.1d). Gains in student confidence were largely unrelated to pre-activity interest in disease ecology, though unlike for the Lotka-Volterra app, students with the least interest in disease ecology appear to have benefitted substantially from the activity (Fig. S2.1e). Broken down by specific concepts, students' understanding of how vaccination rates influence disease dynamics was the only concept substantially different from gains in the control category (Fig. S2.1f).

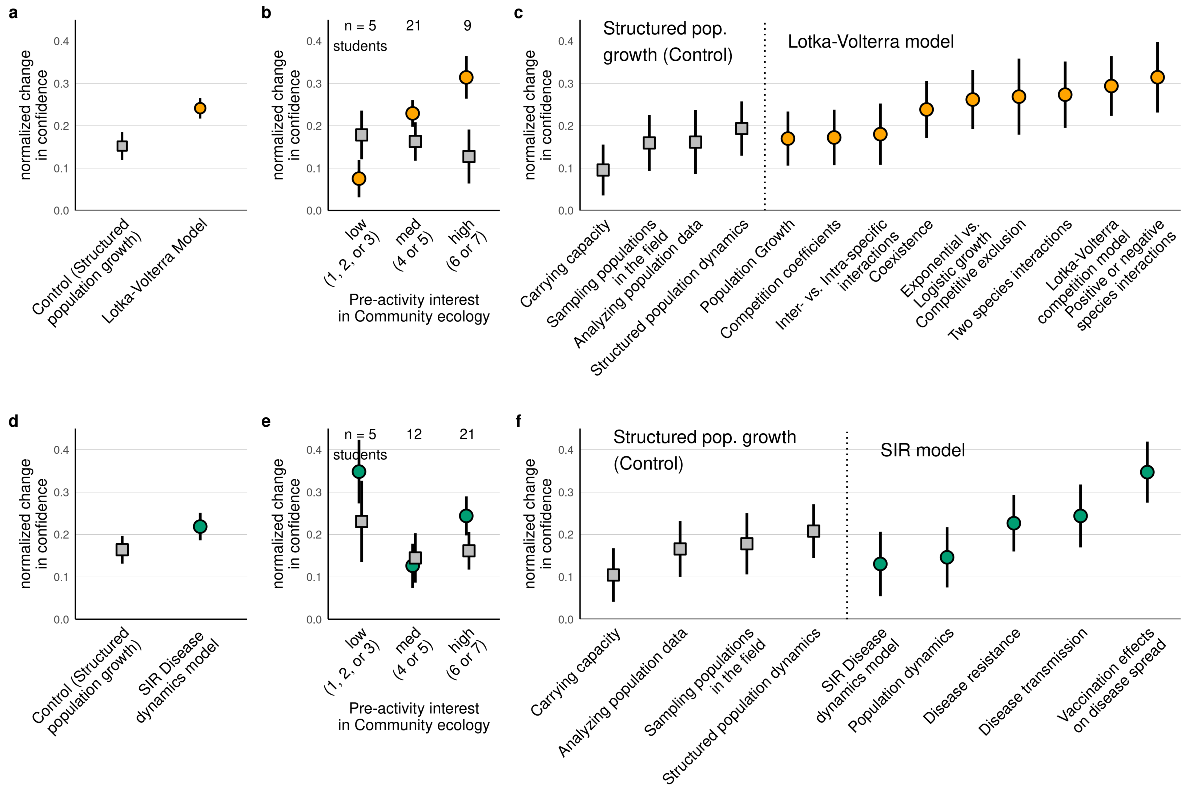

**Fig. S2.1:** Normalized change in MU students' confidence before and after completing a learning activity that incorporated the relevant apps. Panels (a) and (d) show mean and SE (n = 32 and n = 35, respectively) for all concepts relevant to the models covered in the focal apps, or for concepts covered in the "Structured Population Growth" module of the course, which served as controls. Panels (b) and (e) show variation in normalized gains according to students' pre-activity interest in community ecology or disease ecology, respectively. Panels (c) and (f) show normalized gains in specific topics related to the model covered in the focal app or in the control group. summarized across all concepts relevant to each model. In all panels, grey squares indicate questions from the control category (structured population growth), and the orange/green circles indicate questions from the experimental categories.

### Supplement S3: Teaching materials/worksheets used at UCLA and MU

*Supplement S3-1: UCLA extra credit worksheet*

##### I. Pre-Activity Survey (Optional/Anonymous)

##### II. Part 1: Lotka-Volterra Competition Model

- Watch the video on CCLE
- Use Lotka-Volterra competition app: https://ecoevoapps.shinyapps.io/lotka-volterra-competition/
- Using any parameters of your choice, explore these 4 scenarios:
  - Competitive exclusion:
    * K2 < K1/alpha and K1 > K2/beta
    * K2 > K1/alpha and K1 < K2/beta
  - Priority effects/coexistence:
    * K2 > K1/alpha and K1 > K2/beta
    * K2 < K1/alpha and K1 < K2/beta
- For each scenario:
  - Look at the time series graph-What happens to species 1 and species 2 in the long run? Does one of them go extinct? Why?
  - Look at the isoclines and the arrow that shows the combined population trajectory- what happens in each case? What is driving this pattern? Do the outcomes change by using different initial population sizes (while keeping everything else the same)?
- Case Study 1: Honeybees (Species 1) compete for nectar with bumblebees (Species 2) - At a certain habitat patch, the initial density of honeybees is 120 while the initial density of bumblebees is 40. The parameters are listed as follows: $r_1$=0.2, $r_2$= 0.3, $K_1$= 150, $K_2$= 200, alpha=1.2, beta=0.7. Given the initial conditions, what is the expected outcome?
- Case Study 2: Think of two migratory bird populations: species 1 and species 2. The parameters are listed as follows: $r_1$= 0.4, $r_2$ = 0.3, $K_1$= 500, $K_2$= 400, alpha=1.6, beta=1.2. These two species compete every year for nesting sites on cliffsides on the coast of Peru. In 2018, the number of individuals of species 1 that arrived at the beginning of the breeding season was 90, while 150 individuals of species 2 arrived. The following year, 140 individuals of species 1 arrived and 150 of species 2 arrived. Given the initial conditions, what is the expected outcome? What is happening?

##### III. Part 2: Island Biogeography

- Watch the video on CCLE
- Read the introductory text: (optional) https://web.stanford.edu/group/stanfordbirds/text/essays/Island_Biogeography.html
- Use Island Biogeography app: https://ecoevoapps.shinyapps.io/island-biogeography/

- Using any parameters of your choice, explore the 4 cases:
  - Larger island is more diverse than smaller island;
  - Both larger and smaller island are similarly diverse;
  - Smaller island is more diverse than larger island;
  - Both islands are less diverse compared to the initial settings (species pool size).
- Study case:
  - Imagine two similar sized islands, A and B (1 km^2 each), that are 2 and 6 km away from the mainland, respectively; now imagine island B will expand its area at a rate of 1 km^2 every 1,000 years.

**IV. Take the quiz (required for extra credit points)- feel free to use the competition/island biogeography app as you solve the quiz (if needed).**

**V. Post-Activity Survey (Optional/Anonymous)**

*Supplement S3-2: UCLA quiz*

1. What is the outcome of Case Study #1?

    a. Priority effects
    b. **Competitive exclusion of honeybees by bumblebees**
    c. Competitive exclusion of bumblebees by honeybees
    d. Stable coexistence of honeybees and bumblebees

2. What is the outcome of Case Study #2?

    a. Species 1 wins
    b. **Priority effects**
    c. Stable coexistence of species 1 and species 2
    d. Species 2 wins

3. What is the coexistence criterion?

    a. K2 < K1/alpha and K1 > K2/beta
    b. K2 > K1/alpha and K1 < K2/beta
    c. K2 > K1/alpha and K1 > K2/beta
    d. **K2 < K1/alpha and K1 < K2/beta**

4. In other words, for two species to coexist, **intraspecific** competition needs to be greater than **interspecific** competition.

5. What happens in the priority effects scenario? (select all that apply)

    a. Species 1 always wins
    b. **There will be no coexistence**
    c. Species 2 always wins
    d. Competing species coexist
    e. **The outcome of competition depends on the initial conditions**

6. The Lotka-Volterra competition model is useful to show (Select all that apply):

    a. **Under what circumstances can two species coexist**
    b. Predict the outcome of predator-prey interactions
    c. **Predict the outcome of interspecific competition**
    d. Quantify niche overlap

7. Regardless of its size, every island that is farther from the mainland will have:

    a. Larger extinction rates;
    b. Larger immigration rates;
    c. Smaller extinction rates;
    d. **Smaller immigration rates.**

8. For two islands that are equally far from the mainland, the **smaller** one will have a **lower** species richness than the **larger** one, as **extinction** rates decrease with increasing island size, whereas **immigration** rates depend only on the distance from the mainland.

9. Can a large island be less diverse than a small one?

   a. No, the larger island will always have smaller extinction rates, thus higher species richness.
   b. No, the larger island will always have larger immigration rates, thus higher species richness.
   c. **Yes, the larger island will always have smaller extinction rates, but this may be offset by a greater distance from the mainland.**
   d. Yes, the larger island may have greater extinction rates if it is too far from the mainland.

10. What are the species richness of islands A and B, respectively, in the beginning of the study-case scenario (1 km^2 each, 2 and 6 km away from the mainland, respectively)?

    a. $S_A = 33$; $S_B = 14$

11. How long will it take for island B to have as many species as island A?

    a. t = 2,000 years

12. The IB model is useful to show (select all that apply):

    a. **How immigration can be an important driver of species diversity;**
    b. **That species richness may be constant over time, although individual species are coming and going all the time;**
    c. That species whose niche overlaps too much end up being excluded competitively.

*Supplement S3-3: MU Lotka-Volterra competition worksheet*

At Home Lab – Lotka Volterra

This week for your at-home lab we will be using simulations of the Lotka-Volterra equations to better understand how these dynamics work, and the consequences of altering parameters in the equations for competitive dynamics. You will be using this web-based simulation of Lotka- Volterra equations created by our scientist spotlight Dr. Gaurav Kandlikar.

https://ecoevoapps.shinyapps.io/lotka-volterra-competition/ (Links to an external site.)

For this week, you will need to go through the steps outlined below, and write answers to all of the numbered questions. You will need to turn in a document that answers all the questions below. You can complete this assignment alone, or you can work in your groups. If you work in your group, you must submit your own assignment in your own words. If you submit the exact answers as someone else, you both will receive a zero.

**Part 1 - competition coefficients alpha and beta**

First, let’s remind ourselves that Lotka-Volterra is very similar to population growth equations but has the added influence of the other competitor in the equation.

1. What are alpha and beta in the equations? What do they represent?

Leave all of the parameters the same as when you opened the website, and change the alpha and beta parameters to 0.01 (as small as they go).

2. Describe the population dynamics of the two species in this figure? Does this look familiar? What does it look like? Are these two species coexisting under these parameters? What is your evidence?

Now systematically increase and decrease the alpha and beta parameters independently, but leave all other parameters the same.

3. Take a screenshot of the following cases: A) a set of alpha and beta values where species 1 competitively excludes species 2. B) a set of alpha and beta values where species 2 competitively excludes species 1. C) a set of alpha and beta values where both species coexist, but species 2 has a higher population size than species 1 by the end of the simulation (at time 100), and alpha and beta are each greater than 0.4. D) a set of alpha and beta values where both species coexist, but species 1 has a higher population size than species 2 by the end of the simulation (at time 100), and alpha and beta are each greater than 0.4. Include these screenshots in your response, and add figure legends that include your alpha and beta values for each case. What does this tell you about alpha and beta?

**Part 2 - population growth rate**

Hit refresh on the website so the parameter values return to their initial conditions. Now let's explore how growth rates affect the outcome of Lotka-Volterra competition. To isolate growth rate let's set all other parameters to be equal and change growth rates systematically to see their effect.

4. What happens when you set the alpha and beta values to both be 0.3, the carrying capacities for both species to be 200, and the initial population sizes to be 50 for both species? Take a screenshot, put this in your write up, and include a figure legend. What explains the differences in these growth curves?

5. If the carrying capacity for both species is 200, why does the population size for either species never get higher than ~170?

6. Find a situation where one species reaches its carrying capacity. How is this species able to reach this population size?

7. Now, set both K's to be 200, both initial population sizes to be 50, and both competition coefficients (alpha and beta) to be 0.9. What do you see when your r1 (growth rate for species 1) = 0.75, and r2 (growth rate for species 2) = 0.1? Take a screenshot, add a legend, and include it in your writeup. Are these species coexisting? Why do you think this? **Without changing any parameter values before you do this**, make a hypothesis about what will happen to the population dynamics of both species if the simulation runs for 1000 time steps given the parameter values above. Explain why you made this hypothesis.

8. To test your hypothesis, change the length of your simulation to 1000. Take a screenshot of your result, add a figure legend and include it in your writeup. Does this support or refute your hypothesis. Why or why not? If this figure refutes your hypothesis, explain why this outcome occurred.

**Part 3 - Initial Conditions and Carrying Capacity**

Now we will explore how carrying capacity and initial population size alter competition under Lotka-Volterra dynamics. Refresh your website so your parameters return to their initial values.

9. Change the carrying capacity of species 1 (K1) to 50. What population outcome do you see? Take a screenshot, add a legend, and include this in your report.

10. Leaving your carrying capacities at K1 = 50 and K2 = 200, can you find a set of parameter values where Species 1 does not go extinct? Take a screenshot, add a legend, and include it in your write up. Provide all of your parameter values (e.g. alpha = 5). What does it take for species 1 to be able to coexist when it has a very low carrying capacity? How much does initial population size matter relative to the species' growth rates, relative to their competition coefficients alpha and

beta?

*Supplement S3-4: MU SIR worksheet*

**At Home Lab 2 SIR Simulations**

This week we will be exploring SIR models with simulations. While we are off in our units (SIR models were in Unit 2), I believe working through these models will help you understand the concepts better. **You can work with your groups for this assignment if you want, but you need to turn in your own work, in your own words.**

We will be using another simulation created by Dr. Kandlikar, which you can find at this link. https://ecoevoapps.shinyapps.io/sir_disease_dynamics/ (Links to an external site.) We will be focusing on the SIR and SEIR models. Please answer all of the following questions below on a separate word document and submit the answers.

**Part 1 - SIR Models**

Go to the SIR page, and refresh so your parameters are all at their starting values.

**Question 1**: What do the S, I and R parameters stand for in this model? (1 point)

**Question 2**: When your vaccination rate is zero (as it is when you refresh the website), interpret the graph of population size with time. What does this figure tell you?? What does this mean for the potential for the virus to spread in these populations? Why do you think this? (2 points)

**Question 3**: What happens when you set your vaccination rate = 1? Take a screen shot of this new figure, and interpret it (what is it telling you?). What does this mean for the potential for the virus to spread in these new populations? (2 points)

**Question 4**: With the parameters set as they were for question 3, is there any value of the vaccination rate that completely eliminates the potential for the virus to spread in the population? How did you come to this answer? (1 points)

**Question 5**: We have seen that at a low infection rate (0.01), it is difficult to eliminate the potential for virus spread in the population unless 100% of individuals are vaccinated. Refresh the website, and then increase your infection rate to 0.2. Take a screen shot of this new figure, and interpret it (what is it telling you?). Can the virus still spread in the population? Why or why not? How does the population size of the infected individuals explain your answer to if the virus can still spread in the population? (2 points)

**Question 6**: Keeping the parameters the same as for question 5, try to find a vaccination rate that completely eliminates the potential for virus spread. What is it? (1 point)

**Question 7**: Based on your answers and exploration of the SIR models so far, how does increasing infectivity of a virus influence its spread and ease of elimination from a population? (2 points)

**Part 2 – SEIR Models**

Go to the SEIR page, and refresh so your parameters are all at their starting values. This SEIR model is very similar to COVID-19, and how it affects our population.

**Question 8**: What do the S, E, I and R parameters stand for in this model? (1 point)

**Question 9**: Explore the SEIR model by changing parameters and examining outcomes. Based on your exploration, provide 2 things you have learned about the SEIR model. For each, provide your parameter values, and a corresponding figure. Explain what you were investigating, and what insight you gained from the graph (what it is telling you based on the data available). (4 points)

#### Supplement S4: Surveys for assessing teaching outcomes

##### *Supplement S4-1: UCLA survey*

Using interactive apps to teach mathematical models in a remote-learning ecology course

Pre-activity Survey:

I. Select the math classes you have taken:

- Calculus (Math 3A/B/C or LS 30A/B)
- Differential Equations
- Linear Algebra
- Other (write in)

II. From a scale of 1 (not interested) to 7 (very interested), please rate your interest in the following topics:

- Population ecology
- Disease ecology
- Community ecology
- Global change ecology
- Ecosystem ecology
- Conservation ecology

III. From a scale of 1 (not confident) to 7 (very confident), please rate your confidence in the following subjects:

- Exponential vs. Logistic growth
- Carrying capacity
- Population dynamics and time series
- Molecular ecology
- Population growth rates
- Competitive interactions
- Competition coefficients
- Competitive exclusion
- Biogeochemical cycles
- Coexistence
- Lotka-Volterra competition model
- Island Biogeography
- Immigration rate
- Extinction rate
- Mainland/Island Dynamics

Post-activity Survey:

I. From a scale of 1 (not helpful) to 7 (very helpful), how helpful were the interactive apps as a way to generally learn mathematical models in this course?

- Lotka-Volterra Competition
- Island Biogeography

II. From a scale of 1 (not helpful) to 7 (very helpful), please rate how much the interactive apps helped you learn the following concepts (Will also add a not applicable choice):

- Exponential vs. Logistic growth
- Carrying capacity
- Population dynamics and time series
- Molecular ecology
- Population growth rates
- Competitive interactions
- Competition coefficients
- Competitive exclusion
- Biogeochemical cycles
- Coexistence
- Lotka-Volterra competition model
- Island Biogeography
- Immigration rate
- Extinction rate
- Mainland/Island Dynamics

III. What concepts did these model simulations help you understand better? [Free response]

IV. Do you suggest any improvements that would make the apps more valuable as a tool to learn these models? [Free response]

*Supplement S4-2: MU survey*

**Questions after specific webapps:**

From a scale of 1 (not interested) to 7 (very interested), please rate your interest in the following topics:

1. Population ecology
2. Disease ecology
3. Community ecology
4. Global change ecology
5. Ecosystem ecology
6. Conservation ecology

From a scale of 1 (not confident) to 7 (very confident), please rate your confidence in the following subjects:

1. Exponential vs. Logistic growth
2. Carrying capacity
3. Sampling populations in the field
4. Analyzing population data
5. structured population dynamics
6. population growth rate
7. Fecundity effects on population dynamics

From a scale of 1 (not confident) to 7 (very confident), please rate your confidence in the following subjects:

1. Exponential vs. Logistic growth
2. population growth
3. two species interactions
4. competition coefficients
5. positive or negative species interactions
6. interspecific interactions vs intraspecific interactions
7. competitive exclusion
8. coexistence
9. Lotka-Volterra competition model

From a scale of 1 (not confident) to 7 (very confident), please rate your confidence in the following subjects:

1. SIR disease dynamics models
2. disease resistance
3. population dynamics
4. disease transmission

**End of class survey questions**

From a scale of 1 (not helpful) to 7 (very helpful), how helpful were the interactive apps as a way to generally learn mathematical models in this course?

From a scale of 1 (not helpful) to 7 (very helpful), please rate how much the interactive apps helped you learn the following models:

1. Structured population growth
2. SIR disease dynamics
3. Lotka-Volterra competition

What concepts did these model simulations help you understand better? [Free response] [Optional]

Do you suggest any improvements that would make the apps more valuable as a tool to learn these models? [Free response] [Optional]

#### Supplement S5: Student comments on their learning experience with EcoEvoApps at UCLA

##### What concepts did these model simulations help you understand better?

- It was nice to have visual representation of the data. I could physically see how different input values were affecting the results, which was really helpful in making connections.
- Helped clarify the lot Volterra model a lot more
- i liked when there were graphs and each part was explained but i would like an example problem to solidify it
- Conditions that lead to coexistence or competitive exclusion
- Carrying capacity of island biogeography
- Lotka Volterra Model
- I liked the island biogeography model because the equations are difficult to visualize.
- Understanding how to use variables to compare the strength of inter- and intra-specific competition and make predictions with that information. Visualizing how island size and distance affect immigration and extinction rates.
- The stimulations helped to understand both the models because I was able to apply to real life settings
- larger islands and the distance of the island having an impact on extinction rates.
- Both Lotka-Volterra and island biogeography.
- Logistic growth and carrying capacity
- The Lotka-Volterra model was super helpful in understanding the mathematics behind competitive interactions. Similarly, the island biogeography model helped me to better understand immigration, emigration, and interspecific competition
- Population dynamics
- they really helped put all the concepts together because you were able to see how tweaking one thing affects another
- How population dynamics occur through time and then how two populations interact with each other differently through time.
- How changing certain parameters lead to changes in population size
- It helped me understand competitive interactions as well as the impact that initial population numbers have
- Both concepts of Island Biogeography and Lotka-Volterra were better understood after using both apps. I personally feel like I learned more from the island biogeography app more but that could just be a testament to the concept being easier for me. Specifically, I understand the immigration and extinction rates for island biogeography very well. I think more information may be needed for Lotka-Volterra is needed on the app because I learned more from Rosa's video than the app itself.
- Lotka Voltera
- The model simulations helped me best understand the island biogeography theory and the Lotka-Volterra model.

- Lotka Volterra dynamics
- These mode simulations helped me better understand population dynamics.
- I now understand the impact of the competition coefficient on the stability or unstable equilibriums
- The Lotka- Volterra app helped me visualize the concepts learned in lecture, and adjusting the settings and tinkering on my own helped me understand the model better
- The Lotka Volterra model simulation was helpful in putting the difficult math in context for me, although it didn't help me understand the meaning of the math and how to apply it very well. This might just be my fault for having a harder time with mathematical models. The island biogeography model was interesting and I particularly enjoyed the animations. That simulation helped me understand the concept a lot better
- Definitely competitive exclusion/coexistence since it was easier to visualize
- Lotka-Volterra and Island Biogeography theory and how various variable manipulations affected two species and their ecological interaction
- the island biogeography model was very beneficial to understand the connection between size and distance
- These simulations helped myself better understand Lotka-Volterra Competition Models and competitive exclusion.
- Understanding under which conditions populations coexist and when one overcomplete the other
- They helped me visualize the relationships between each variable using graphs.
- These simulations helped clearly show how Lotka-Volterra is utilized to predict species competition as well as island biogeography theory is used to predict species richness for island- mainland dynamics.
- The Lokta Volterra Model explanation was very helpful!! The Island biogeopgraphy model helped outline the main concepts.
- Island biogeography (specifically how island size and distance each contributes to population dynamics) and Lotka-Volterra equations (specifically what the variables meant)
- Effect of distance versus area for island biography in a way easy to visualize
- competitive interactions between different species and island biogeography concepts
- Population dynamics using island biogeography model
- It really helped with Island biogeography theory! It was really helpful seeing how island sizes and distance affect the model
- These models helped me to better understand competitive interactions between 2 species as simulated by the Lotka-Volterra model, as well as how immigration and extinction rates are affected by the size and distance of islands.
- LV model = Competitive competition, Coexistence, and unstable equilibrium
  Island model = island biogeography model
- growth rates
- The lotka-Voltera simulation helped me understand what the lotka-Voltera predicts because it was more hands on than listening to its explanation during lecture.The

island simulation also made it easy to understand how different variables and values of size/distance affect island populations. Visualizing these concepts made the model very clear.
- These models helped me to better understand competitive interactions and changes in population dynamics over time
- extinction rate
- The model simulations really helped me understand the relationships between the extinction rates and immigration rates and distance in the island biogeography model, but the Lotka- Volterra didn't help me as much, but it still helped me understand the way Lotka-Volterra functions and the different outcomes that it can have.
- How size and distance from the island can impact extinction/immigration rates
- population dynamics and growth rates
- Interspecific competition, immigration, and extinction rates.

**Do you suggest any improvements that would make the apps more valuable as a tool to learn these models?**

- It would be nice to have a sample example given, so that students are able to correctly interpret the results shown on the graphs
- NA, thanks!
- showing more examples and step by step
- not right now
- Not really, both were very specific already
- not that I can think of!
- The lotka voltera interactive is already very helpful, but a blurb in the app about what each of the values means could drive the point home even better. I had to check a few times to make sure my understanding of each variable was correct.The apps were very easy to use. The instructions that Rosa and Marcel provided were excellent for doing the extra credit. I really enjoyed how I can apply real life concepts to the apps
- None. Overall great experience.
- No. It just constantly had to be reloaded if you went off the website for a short while, but that's not really an issue.
- It should be a bit more clearing with its instructions on how to use
- I think that the island biogeography model was almost perfect as is. It was very user friendly and easy to use. The Lotka-Volterra model could be simplified somewhat by reducing the number of variables that can be changed at once, or perhaps by designing sub modules to teach individual concepts
- I enjoyed it!
- sometimes the website would crash and i wasn't able to use itt on the laptop
- The coefficients and all the input values just become numbers after you throw so many at once. It would be helpful to accompany a slow-pace conceptual backing when tweaking each value individually while looking at the model.
- none at this time

- No suggestions!
- I don't have improvements for the island biogeography app; I found it very helpful. An improvement I have for the lotka-Volterra app is addressing what the open and closed dots mean as well as increasing the size of the graphs when the isoclines are intercepted in a closer range. It got really hard to see some effects when the parameters made the isoclines fairly close.
- no, i thought they were good as they were
- The apps were great and truly helped with improving my understanding of certain concepts. The only issue I had was that the website wouldn't work/load at times. I am not exactly sure why, but I would have to refresh the page or visit it at a different time for it to completely load.
- N/A, these models were very helpful. However, just general stability could be improved (the models frequently crashed and needed to be reloaded)
- No
- Provide many questions but also add the answers so you can see why you got a problem incorrect
- It would be helpful if the lines in the island biogeography app are thicker, so they are easier to see with the graphs
- The Lotka Volterra model simulation would benefit from some explanation of how changes to the parameters affected the model. I think there were just a lot of parameters to focus on that it confused me
- It can explicitly say which species won
- n/a; very helpful tools!
- i think it would be beneficial to have the lotka voltera model explained with examples
- I took me a while to understand the results so if the app states "the population will coexist" "population 1 will outcompete population 2," etc. that would be helpful
- The site is a little buggy/laggy.
- Allowing a wider range of values for the parameters to see extreme results.
- The apps worked great. The lotka volterra model simulation is not as user friendly. It is hard to exactly understand which K1>k2/alpha you are inputting. I didn't know how to explicitly tell what I was doing.
- Apps are already super easy to use! Pairing them with the scenarios made it even clearer what the app was trying to demonstrate
- The Lotka-Volterra model would not load for the longest time. It took me around 45 minutes of opening it on different and devices to get it to open and/or stop freezing. I would suggest fixing this app before using it as an educational tool.
- Yes, the Lotka-Volterra app got disconnected a lot. It sometimes stops working and that's discouraging, but it was simple to use overall
- One aspect of the model that can be better explained is the model for coexistence for Lotka-voltera model.
- Maybe you could give some examples at the bottom for people to test before putting their own data in
- I had trouble using the Lotka-Volterra model app as it would freeze periodically and I would be unable to reload the page which made taking the extra credit quiz a

little difficult.

- LV model = include things like K1, K2, etc. to label the lines;Island model = nothing – it's really good
- showing how to use it
- I would suggest ability to hover with the mouse over the variable you are changing, and a pop up box appears explaining clearly how adjusting that variable will affect the results.
- I would suggest adding a key that explains each of the variables that can be manipulated and expanding on how they affect the overall model.
- I think it is helpful
- The Lotka-Volterra models could be a bit more clear in how these different variables affect the outcomes of the models, but the island biogeography one was very clear.
- None, both apps were clearly explained well by both TAs and there was nothing too confusing about them. If anything, they helped me better understand the four possible outcomes of the Lotka-Volterra model better than I did before.
- It looks great
- I think they are pretty good overall. Maybe it'd be better if the instructions on how to use the apps could be clearer
